# A new method to identify global targets of RNA-binding proteins in plants

**DOI:** 10.1101/2021.06.11.448000

**Authors:** You-Liang Cheng, Hsin-Yu Hsieh, Shih-Long Tu

## Abstract

**Background:** RNA-binding proteins (RBPs) play crucial roles in various aspects of post-transcriptional gene expression; their functions can vary between tissues, cell types, developmental stages, and environmental conditions. Identifying RBP target RNAs and investigating whether they are differentially bound by RBPs in different cell types, stages, or conditions could shed light on RBP functions. Although several strategies have been designed to identify RBP targets, they involve complicated biochemical steps and require large quantities of material, and only a few studies using these techniques have been performed in plants. The TRIBE (targets of RNA binding proteins identified by editing) method was recently developed to identify RBP targets using a RBP coupled to the catalytic domain of a *Drosophila* RNA editing enzyme and expressing this fusion protein *in vivo*. The resulting novel editing events can be identified by sequencing. This technique uses little material and does not require complex biochemical steps, however it is not yet adapted for use in plants.

**Results:** We successfully applied an optimized genome-wide TRIBE method in plants. We selected the splicing regulator polypyrimidine tract-binding protein (PTB) as a model protein for testing the TRIBE system in the moss *Physcomitrium patens*. We demonstrated that 13.81% of protein-coding gene transcripts in *P. patens* are targets of PTB. Most potential PTB binding sites are located in coding sequences and 3’ untranslated regions, suggesting that PTB performs multiple functions besides pre-mRNA splicing in this moss. In addition, TRIBE showed reproducible results compared to other methods.

**Conclusions:** We have developed an alternative method based on the TRIBE system to identify RBP targets in plants globally, and we provide guidance here for its application in plants.

## Background

RNA-binding proteins (RBPs) are crucial for many post-transcriptional steps of gene expression, such as precursor messenger RNA (pre-mRNA) splicing, RNA processing, and the modification, nuclear export, and translation of mRNA (1-4). The ability of RBPs to associate with RNA is essential for their functions. RBPs are divided into different groups with diverse functions based on their binding affinity and selectivity in recognizing specific target RNA sequences and structures. In addition, the same RBP may bind to distinct RNA transcripts in different cell types, tissues, developmental stages, or environmental conditions. Therefore, identifying the targets of RBPs is an important task for revealing the physiological functions of these proteins (5-7).

RNA immunoprecipitation (RIP) and crosslinking immunoprecipitation (CLIP) have been used to identify the targets of RBPs (8). Both methods rely on crosslinking and the use of antibodies for immunoprecipitation. An RNA sample is first treated to covalently link RBPs with their RNA targets and then fragmentated by sonication or nuclease digestion. The remaining RBP-bound RNA is immunoprecipitated using a specific antibody, eluted from an antibody-conjugated matrix, and used to construct libraries for RNA sequencing (RNA-seq) (9,10). The complicated biochemical steps required in these methods increase the difficulty of detecting RBP targets, especially when comparing data between samples. In addition, due to the low (∼1-5%) efficiency of crosslinking by UV light during CLIP, only a small amount of RBP-bound RNA can be detected, posing fundamental problems including a low signal-to-noise ratio and high rate of false positive results. Furthermore, the inefficiency of antibody-based biochemical methods necessitates large amount of starting material, making it difficult to study RBP targets in single cells or tissue types.

A method called TRIBE (targets of RNA binding proteins identified by editing) was recently established to identify RBP targets in *Drosophila melanogaster* (11). The TRIBE method takes advantage of the catalytic domain of the *Drosophila* enzyme ADAR (adenosine deaminase acting on RNA), which converts adenosine to inosine (A-to-I conversion) in its target RNAs. The RBP of interest is fused to the catalytic domain of ADAR (ADARcd). When expressed in cells, the RBP-ADARcd fusion protein edits its RNA targets, resulting in novel editing events. Total RNA extracted from the cells can be subjected to Sanger sequencing or RNA-seq to further identify these editing sites, which represent potential RBP targets. The TRIBE method was recently further updated to the hyperactive TRIBE (HyperTRIBE) method. This technique employs a hyperactive ADAR harboring a substitution of glutamate 488 by glutamine, which increases the enzyme’s sensitivity and reduces sequence bias (12,13). This powerful method can identify RNA targets of the RBPs of interest using a small quantity of material without the need for complicated biochemical processes. Compared to RIP and CLIP, TRIBE allows RBP targets to be identified in a cell-type- or tissue-specific manner and to compare RNA binding activity between samples.

Pre-mRNA splicing is regulated by various splicing factors, such as serine/arginine-rich (SR) proteins and heterogeneous nuclear ribonucleoproteins (hnRNPs). Early studies indicated that SR proteins function as positive regulators of pre-mRNA splicing and hnRNPs function as negative regulators of this process (14). However, splicing regulation appears to be a more complicated process due to the combinations of splicing factors and regulatory RNA elements identified (15,16). Polypyrimidine tract-binding protein (PTB) is a well-studied and evolutionarily conserved hnRNP that binds to the polypyrimidine tracts following the branch-point adenosine on intronic regions of pre-mRNA (17). Several mechanisms have been proposed for PTB-mediated intron splicing (18). In *Arabidopsis thaliana*, three genes encoding PTB1, PTB2, and PTB3 have been identified, with PTB1 sharing high sequence similarity with PTB2 but not with PTB3. Although PTBs have been shown to be involved in pollen development and abscisic-acid-dependent seed germination, the detailed mechanisms have not been addressed (19,20). Plants with defective PTBs show differential gene expression and alternative splicing of various genes (19). However, whether these gene transcripts are direct targets of PTBs is still unknown. TRIBE accompanied by RNA-seq could provide more detailed information about the regulation of PTBs.

In the current study, we successfully established the HyperTRIBE method for use in plants. Using HyperTRIBE in the moss *Physcomitrium patens* (*P. patens*), we determined that 13.81% of protein-coded genes were successfully edited by PTB2. Most editing sites were located in coding sequences (CDS) and 3’ untranslated regions (3’ UTRs), suggesting that PTB might have multiple functions other than pre-mRNA splicing. In addition, some editing sites were located near or within RIP-enriched regions, suggesting they are potential binding regions of RBP. Overall, we demonstrated that the HyperTRIBE system is functional in plants and is a simple method for identifying RBP targets.

## Results

### The HyperTRIBE pipeline

To determine whether the HyperTRIBE system could be used in plants, we optimized the coding sequence of *Drosophila ADARcd* for plant expression. We introduced a point mutation in the active site of the protein to generate a hyperactive version of ADARcd (hereafter referred to as HyperADARcd) (21,22). A hnRNP protein homologous to Arabidopsis PTB2 (AT5G53180) in *P. patens* was chosen for analysis because PTB is a relatively well-studied RBP that directly binds to RNA. We constructed *pPGX8* plasmids encoding *PTB2-HyperADARcd-2myc* and *HyperADARcd-2myc* (hereafter called *PTB2-HyperTRIBE* and *HyperADARcd*, respectively) and used them to transform *P. patens* protonemal cells. Both fusion proteins were expressed under the control of a β-estradiol-inducible promoter. We measured the levels of PTB2-HyperTRIBE and HyperADARcd in transformed protonemal cells by immunoblotting (Figure S1), extracted total RNA from two biological replicates, and used the RNA to prepare libraries for RNA-seq (Figure 1).We mapped the sequence reads to *P. patens* genome V3.3 using the Bowtie2 and BLAT programs (23,24) and counted the nucleotides at each position of the genes using the PolymorphismCounter and NTFreqCounter functions in the RACKJ package. We defined an editing site as a base position with an adenosine-to-guanosine (A-to-G) conversion in *PTB2-HyperTRIBE*-overexpressing cells compared to the same position in wild-type cells, with significant differences detected by a *G*-test. We assigned the editing sites to the corresponding genes by gene annotation and subjected the sites that were identified in both biological replicates to further analysis. The detailed criteria used are described in the Methods.

**Figure 1.**
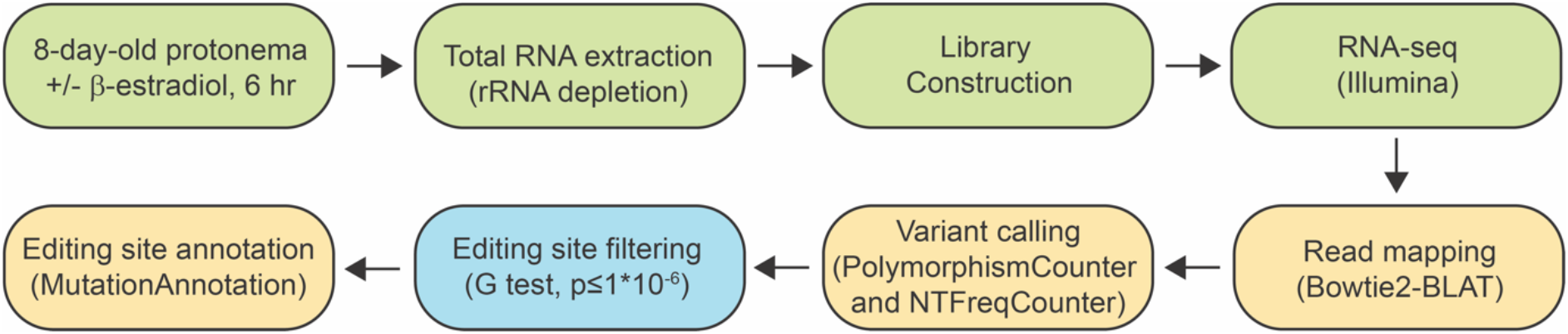
HyperTRIBE workflow. Total RNA was extracted from protonemata and subjected to RNA-seq. The reads were mapped to the reference genome using Bowtie2 and BLAT, and the number of A-to-G conversions in each gene was counted using the PolymorphismCounter and NTFreqCounter functions of the RACKJ package. Editing sites defined based on statistical significance, as determined by *G*-test, were further analyzed.

### HyperTRIBE analysis to identify PTB2 targets

The number of editing sites (*n* = 13128) increased dramatically after PTB2-HyperTRIBE induction (Figure 2A), suggesting that PTB2-HyperTRIBE indeed catalyzes adenosine deamination in moss cells. The expression of the inducible promoter was well controlled, as editing sites (*n* = 97) were rarely detected in *PTB2-HyperTRIBE* cells in the absence of β-estradiol (Figure 2A). A comparison of the editing levels at the same site in two replicates showed that the editing events were reproducible (*R*^2^ = 0.86) (Figure 2B), indicating that PTB2-HyperTRIBE has specificity toward its substrates. Overall, the editing levels at the same sites were higher in replicate 2 than in replicate 1 (Figure 2B); this difference might have been due to a batch effect. RNA was also edited in *HyperADARcd* cells (Figure 2A). Approximately 29.83% (3017 out of 10,111) of editing sites were detected in both *PTB2-HyperTRIBE* and *HyperADARcd* cells (Figure 2C). We randomly selected 100 editing sites from overlapping regions and found that most showed higher levels of editing in *PTB2-HyperTRIBE* cells than in *HyperADARcd* cells (Figure 2D). These results suggest that non-specific editing by HyperADARcd could occur, but conjugated RBP still showed RNA binding affinity and specificity.

**Figure 2.**
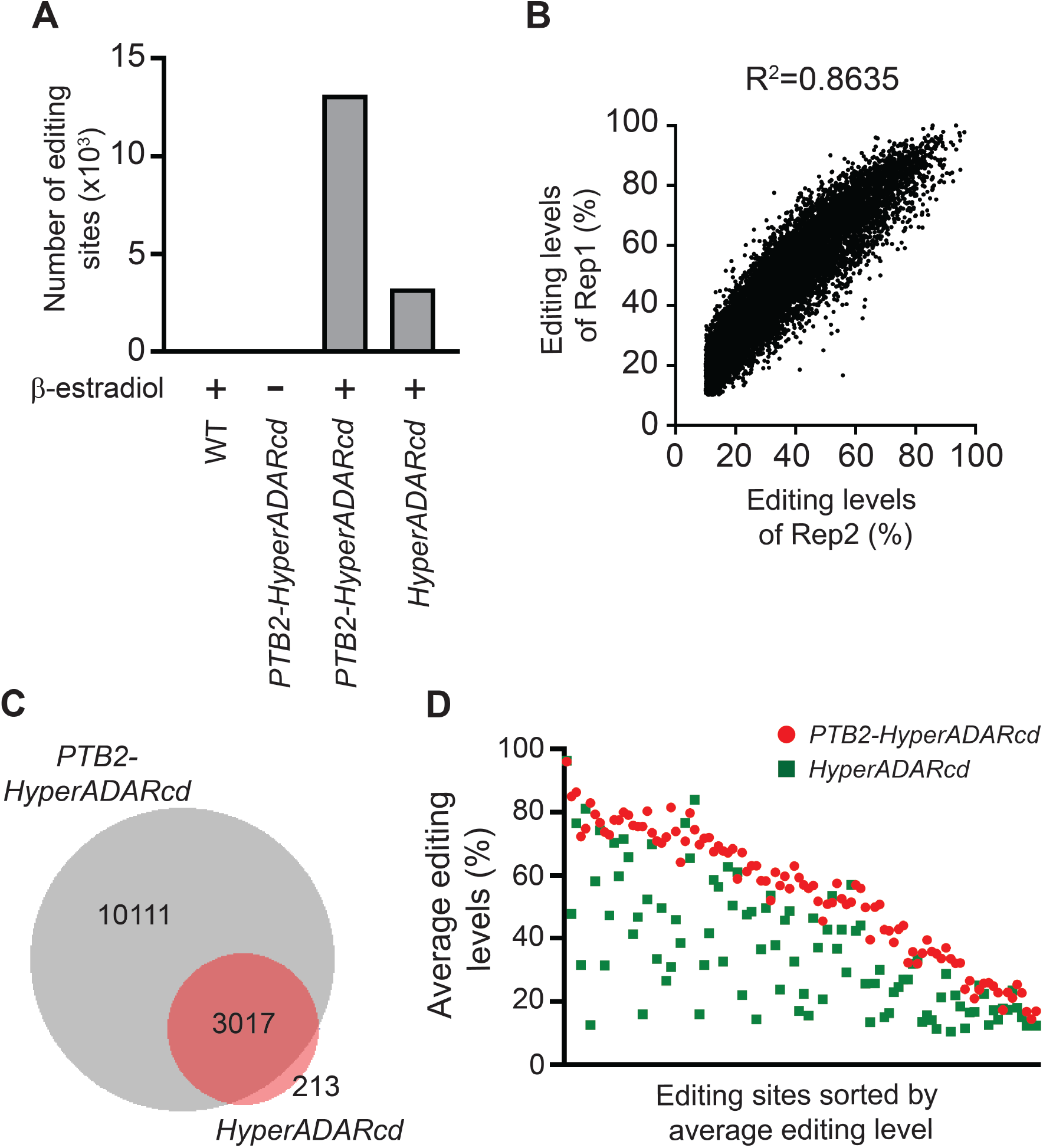
HyperTRIBE analysis in different cells. (A) Number of editing sites in wild-type (WT), *PTB2-HyperTRIBE*, and *HyperADARcd* cells in the absence or presence of β-estradiol. The number of A-to-G conversions increased in *PTB2-HyperTRIBE* cells after β-estradiol induction but were rarely observed in the absence of β-estradiol. (B) The correlation of editing levels in two biological replicates (designated Rep1 and Rep2). (C) Venn diagram comparing the editing sites in *PTB2-HyperTRIBE* and *HyperADARcd* cells. (D) Comparison of editing levels in 100 randomly chosen editing sites from an overlapping region (*n* = 3017, Fig. 2C) in *PTB2-HyperTRIBE* and *HyperADARcd* cells. Most of the sites show higher editing levels in *PTB2-HyperTRIBE* cells than in *HyperADARcd* cells.

To increase the stringency of HyperTRIBE, we removed the 3017 editing sites from the list of *PTB2-HyperTRIBE* cells. In total, 7094 editing sites were obtained and further annotated. These editing sites resided in 4546 RNA transcripts, comprising approximately 13.81% of protein-coding genes in *P. patens*, confirming the notion that PTB is a general splicing regulator associated with many mRNAs. In addition, 48.31% of PTB-targeted transcripts contained only one editing site (Figure 3A). The number of editing sites did not increase proportionally with increasing transcript abundance or length (Figure S2), indicating that PTB2-HyperTRIBE has substrate specificity. In addition, most editing sites resided in CDS and 3’ UTRs (Figure 3B). These results suggest that PTB2 in *P. patens* has diverse roles in post-transcriptional regulation, such as pre-mRNA splicing, RNA stability, and polyadenylation.

**Figure 3.**
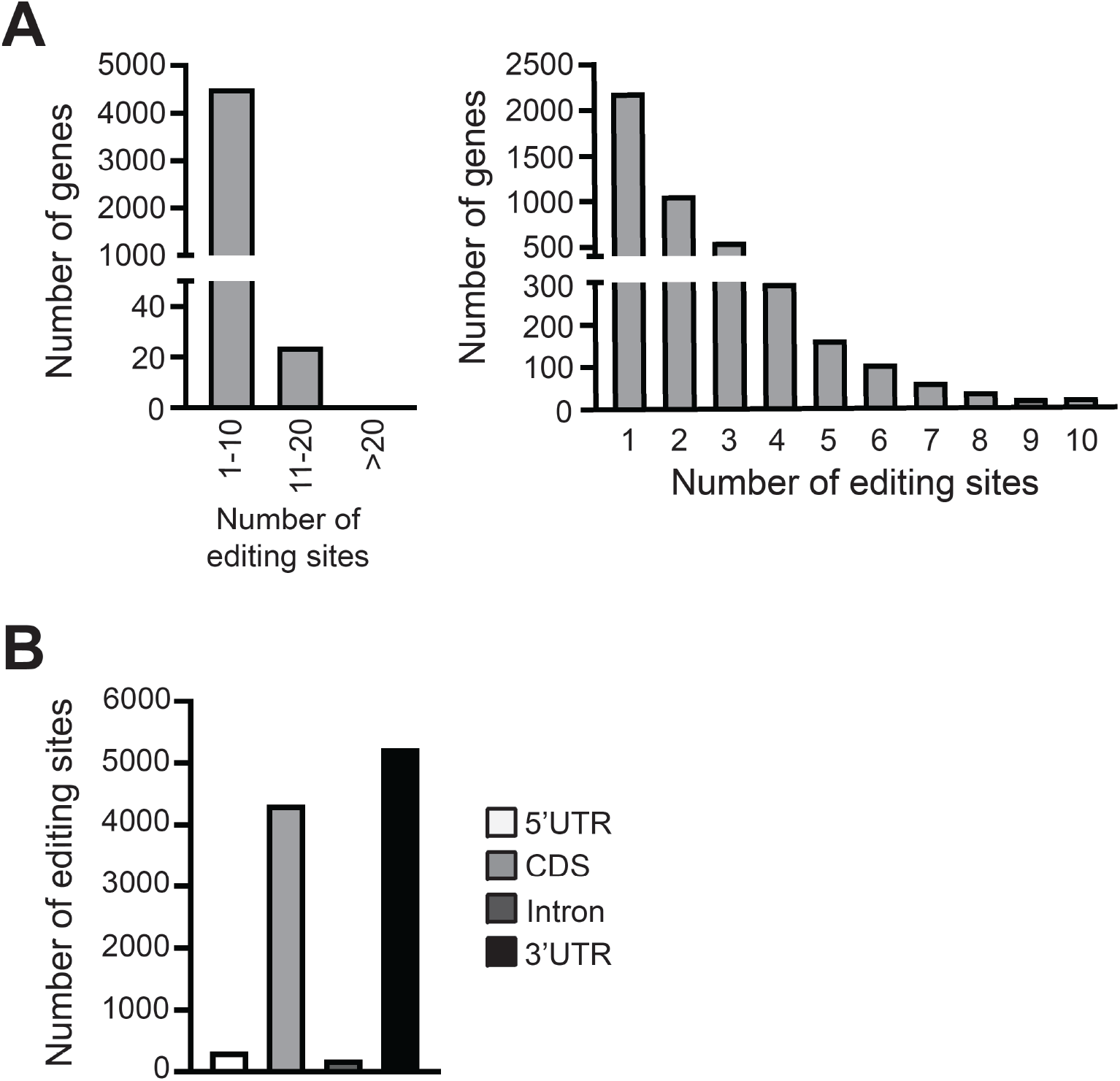
Number and distribution of editing sites in genes. (A) Histogram of the number of editing sites per gene. The group with 1-10 edits per gene was further divided into ten categories, shown at right. The majority of genes contained one or two editing sites. (B) Distribution of the number of editing sites in different types of gene regions. Most of the editing sites resided in CDS and 3’ UTRs.

### Editing sites in HyperTRIBE are located near or within RIP-enriched regions

Because the RNA targets of PTB in plants are unknown, we used another genome-wide method, RNA immunoprecipitation sequencing (RIP-seq), to confirm the PTB2 targets identified by HyperTRIBE. We generated a PTB2-5myc-overexpressing *P. patens* line for RIP-seq. We chose PTB2-5myc instead of PTB2-HyperTRIBE for RIP-seq to reduce the identification of false positive targets because HyperADARcd alone could bind to RNA non-specifically. For RIP-seq, we fixed 8-day-old protonemata expressing PTB2-5myc with 1% formaldehyde for protein-RNA crosslinking, isolated and lysed the nuclei, immunoprecipitated PTB2-5myc using anti-cMyc antibody, and detected this protein by immunoblotting (Figure 4A). We subjected RNA extracted from the immunoprecipitated fraction to RNA-seq, finding that 3494 gene transcripts were enriched in *PTB2-5myc* cells (Figure 4B). Approximately 30.51% of PTB2 targets identified by RIP-seq were also present in the HyperTRIBE data (Figure 4B). For RNA targets identified by both methods, editing sites identified by HyperTRIBE were near or within the enriched regions identified by RIP-seq (Figure 4C). This observation is consistent with the finding that the majority of editing sites identified by HyperTRIBE are located within 500 bp from CLIP peaks (11).

**Figure 4.**
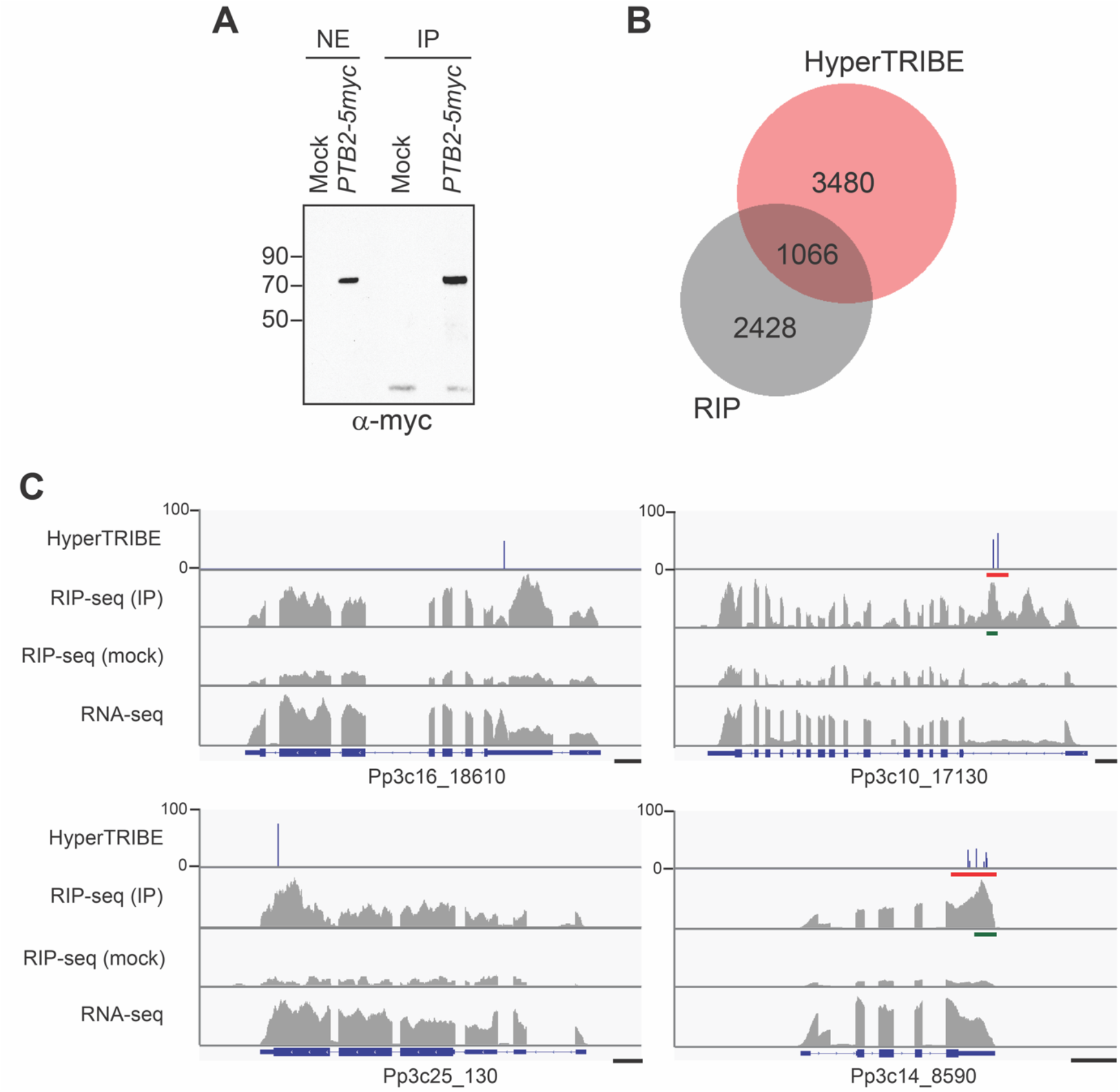
Comparison of the targets identified by RIP-seq and HyperTRIBE analyses. (A) Immunoblot of PTB2-5myc after immunoprecipitation using anti-cMyc antibody. NE, nuclear extract; IP, PTB2-5myc-immunoprecipitated fraction. (B) Venn diagram comparing PTB2 targets identified by HyperTRIBE and RIP-seq. (C) Integrative Genomics Viewer (IGV) plots visualizing PTB2-identified targets by HyperTRIBE and RIP-seq. Editing sites and editing levels from HyperTRIBE and enriched regions identified by RIP-seq and RNA-seq are shown. The editing sites resided near the enriched regions. Scale bars, 500 bp.

### Validation of PTB2 targets

To confirm the HyperTRIBE and RIP-seq results, we ligated ∼500 bp regions of PCR amplicons from target cDNA fragments (3’ UTR of Pp3c14_8590 and intron of Pp3c10_17130) into the *pBlueScript* vector and subjected 27-30 randomly chosen resulting clones to Sanger sequencing. The positions (editing sites) identified by Sanger sequencing and HyperTRIBE were reproducible (Figure 5A), indicating that PTB2-HyperTRIBE exhibits specificity. Enriched fragments identified by RIP-seq were located near the editing sites identified by HyperTRIBE (Figure 5B), which is consistent with our results from genome-wide experiments.

**Figure 5.**
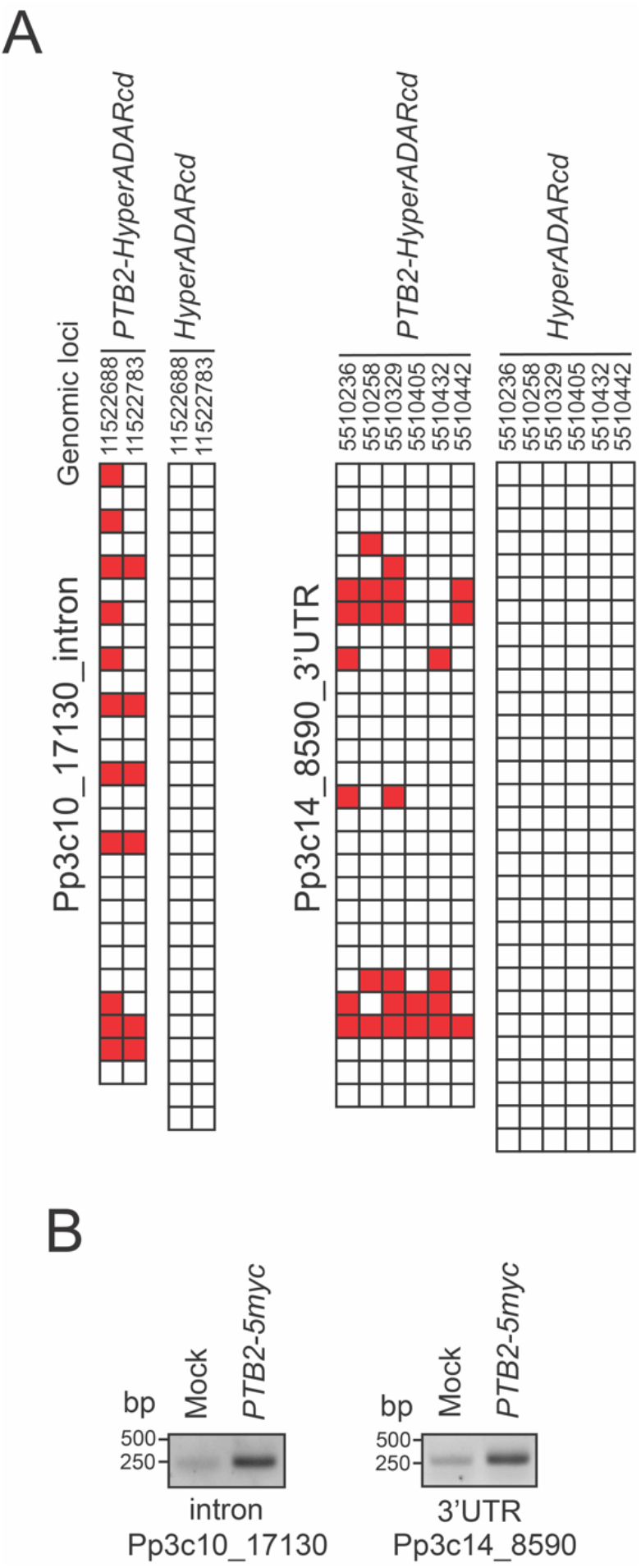
Validation of editing sites and levels in HyperTRIBE and RIP-seq data. (A) PCR amplicons from target cDNAs (red regions in Figure 4C) were ligated into vectors, and 27-30 clones were subjected to Sanger sequencing. The presence of G or A in an editing site is indicated by red or white, respectively. The editing sites identified by both Sanger sequencing and HyperTRIBE were reproducible. (B) RT-PCR was performed after RIP to confirm the enriched region (green bars in Figure 4C) identified by RIP-seq.

## Discussion

Post-transcriptional regulation of gene expression is mediated by a wide range of protein factors that bind to RNA. These RBPs play crucial roles in various aspects of mRNA maturation, especially pre-mRNA splicing. At least 196 RNA-recognition-motif-containing proteins and 26 K homology (KH) domain proteins are encoded in the Arabidopsis genome, but most have not yet been studied (6). RBP activity can vary in different tissues, cell types, developmental stages, and environmental conditions. Uncovering the RNA targets of RBPs would help uncover the physiological roles of RBPs in cells. In the current study, we established the HyperTRIBE platform for plant systems using *P. patens* as a model. This powerful method can reveal the RNA targets of RBPs of interest in specific tissues, cell types, developmental stages, and environmental conditions using a small number of cells without the need for complicated biochemical processes. By combining HyperTRIBE with next-generation sequencing (NGS), we also designed an analysis pipeline that allows us to identify mRNAs associated with RBPs on a large scale.

### Development of the HyperTRIBE method in plants

The TRIBE method was first developed using the RNA-editing enzyme ADAR from *Drosophila* (11). After fusing the catalytic domain of ADAR (ADARcd) with the RBP and expressing this fusion protein *in vivo*, the editing sites in RNA are identified based on the specificity of ADARcd for the UAG sequence in the RBP targets. An updated version of this technique, HyperTRIBE, was developed by substituting glutamate 488 in ADAR with glutamine to reduce the preference of ADARcd for UAG and enhance its deaminase activity so as to increase the chances of identifying specific RNA targets of RBPs of interest (12,13). However, *Drosophila* ADARcd shows low editing efficiency in human PC3 cells (25). To utilize HyperTRIBE in human cells, the catalytic domain of human ADAR2 was used to improve its editing efficiency. In the current study, we optimized the codons of *Drosophila ADARcd* for expression in plants. Our results confirm the notion that *Drosophila* HyperADARcd is stably expressed and functional in moss cells, even when fused with another protein.

Although the mutation of ADARcd reduces the preference for UAG in RBP targets, HyperADARcd alone still performed non-specific editing of RNA in moss cells. This type of non-specific editing was not observed in fruit fly or human cells (13,25). It is possible that human and *Drosophila* cells have regulatory mechanism to better control ADAR activity. Alternatively, perhaps this difference was due to the different subcellular environments in animal and plant cells. The number of editing sites and the editing levels in *P. patens* cells rose in proportion to the length of HyperADARcd induction, increasing the difficulty of identifying the specific targets of RBP-HyperTRIBE. It is possible that this occurred because plant cells are usually occupied by a large vacuole, which reduces the volume of the cytoplasm and increases the chances for non-specific interactions between HyperADARcd and RNAs. To account for this problem, we adopted two strategies. First, we confirmed that the editing sites and editing levels were reproducible between biological replicates. Second, by precisely controlling the expression levels of both PTB2-HyperTRIBE and HyperADARcd, we were able to dissect PTB2-specific mRNA targets from non-specific editing sites. We therefore suggest that to reduce the identification of false positive targets, HyperADARcd should always be used as a control in HyperTRIBE experiments in plant systems. In addition, the induction of RBP-HyperTRIBE and HyperADARcd expression should be well controlled to ensure the same or similar expression levels.

Overexpressing human ADAR2 reduced the growth rate of budding yeast cells, suggesting that the deaminase may edit some RNA transcripts that are important for the physiological functions of cells (26). Therefore, the deaminase activity should be well controlled in cells overexpressing RBP-HyperTRIBE and HyperADARcd. In the current study, we used an inducible promoter to control RBP-HyperTRIBE and HyperADARcd expression to avoid any unexpected effects of these proteins in moss cells. Indeed, very few editing sites were identified when the inducible promoter was turned off. We suggest that protein expression should be checked prior to induction to avoid background expression of the fusion protein due to promoter leakage.

### Data analysis and comparison

We also performed RIP-seq to further confirm the RNA targets identified by HyperTRIBE. Approximately 30.51% of the RNA targets identified by RIP-seq were also detected by HyperTRIBE. Perfect overlap between the two techniques was not expected because they use distinct methods and analysis platforms to identify RBP targets. In addition, these two methods have their own false positive targets. For RIP, non-specific crosslinking and antibody binding cause false positive results. In the case of HyperTRIBE, RBP-HyperTRIBE may also non-specifically edit RNA targets due to the editing preference of HyperADARcd. However, our results indicate that the number of editing sites is not correlated with the length or abundance of transcripts (Figure S2), suggesting that RBP-HyperTRIBE still has specificity toward its substrates. Although some editing sites were located near RIP peaks, neither method has the ability to identify the binding sites of RBPs, which must be determined using other methods, such as CLIP.

### Proposed functions of *P. patens* PTB2

In this study, we selected PTB as the RBP for HyperTRIBE because PTB is a well-studied hnRNP that is evolutionarily conserved in different organisms. Although PTB is reported to bind to the polypyrimidine tracts following the branch-point adenosine in the intronic regions of its targets, we found that most of the editing sites were located in CDS and 3’ UTRs. There are several possible explanations for this observation. First, we prepared total RNA but not nuclear RNA for sequencing. Intronic regions in pre-mRNAs might be edited but spliced out before RNA extraction. Nascent RNA-seq is probably a better choice for identifying the binding targets of RBP to the intronic regions of pre-mRNAs. Second, introns in plants are too short to be edited after RBP binding. It is still possible that PTB2-HyperTRIBE binds to intronic regions and edits nearby CDS or 3’ UTRs. On the other hand, we determined that transcripts encoded by 13.81% of *P. patens* genes were edited by PTB2-HyperTRIBE but not by HyperADARcd, suggesting that PTB2 binds to a broad group of pre-mRNA species for splicing regulation. We also observed that PTB2-HyperTRIBE intensively edits 3’ UTRs, suggesting that *P. patens* PTB2 might have multiple functions other than pre-mRNA splicing, such as mRNA polyadenylation, trafficking, and stability. Further investigation will be required to identify the roles of PTBs in plants.

### Perspectives

The RIP and CLIP methods are imperfect and are not suitable for examining small amounts of material. The TRIBE method employs ADARcd fused with RBP to overcome these issues. TRIBE was further modified to HyperTRIBE to increase its sensitivity and enable the identification of more RBP targets. The method makes it possible to identify RBP targets in a tissue- and cell-type-specific manner and to compare different samples. For example, by combining HyperTRIBE with laser microdissection technology, a single-cell RBPomics technique could be established in the future. In addition, the positive correlation between editing levels and the amounts or binding strength of RBP-HyperTRIBE should be further addressed. If this correlation is indeed present, differential editing events could be used to quantify the interactions between RBP and its targets. Thus, changes in editing levels might represent the dynamics of RBP-RNA interactions. By taking advantage of HyperTRIBE, additional roles of RBPs can be uncovered in future.

## Methods

### Plant growth, transformation, and treatment

*P. patens* protonemata were grown on cellophane-overlaid BCDAT medium (1 mM MgSO_4_, 1.84 mM KH_2_PO_4_, pH 6.5, 10 mM KNO_3_, 45 μM FeSO_4_, 1 mM CaCl_2_, 5 mM ammonium tartrate, 1Δ Hunter’s micronutrients, and 0.8% agar) under continuous white light at 24°C for 7 to 8 days. PEG transformation was performed according to a previous protocol (27). The transformants were checked by immunoblotting. For PTB2-HyperTRIBE induction, 15 ml of 10 μM β-estradiol was added to protonemata at under continuous white light at 24°C for 6 h, and protein expression was confirmed by immunoblotting.

### Plasmid construction

For the inducible construct, *pPGX8-PTB2-HyperADARcd-2myc* was generated. *Drosophila HyperADARcd* (*ADARcd*^*E488Q*^) was codon-optimized, synthesized, and cloned into *pBlueScript-2myc* at the *Eco*RI and *Hin*dIII sites. The coding region of *P. patens PTB2* was cloned into *pBlueScript-HyperADARcd-2myc* at the *Bam*HI and *Eco*RI sites. *PTB2-HyperADARcd-2myc* was further amplified by PCR and cloned into *pENTR* at the *Not*I and *Asc*I sites. Subsequently, *PTB2-HyperADARcd-2myc* from *pENTR* was introduced into *pPGX8* through Gateway cloning.

### Total RNA extraction

Total RNA was extracted from the samples using a GeneMark kit according to the manufacturer’s protocol (GMbiolab, Taiwan).

### Library construction and RNA sequencing

To prepare strand-specific RNA-seq libraries, 4 μg of purified RNA was depleted of ribosomal RNA using a Ribo-Zero rRNA Removal Kit - Plant Leaf (Epicentre) following the manufacturer’s instructions. Libraries for RNA-seq were prepared with an Illumina TruSeq stranded mRNA sample preparation kit (Illumina, USA) according to the manufacturer’s protocol. Briefly, the rRNA-depleted RNA was fragmented and used as a template to synthesize first-strand cDNA with SuperScript III reverse transcriptase (Invitrogen, USA) using dNTPs and random primers. Second-strand cDNA was generated using dUTP mix. A single ‘A’ base was added to the 3’ end of the double-stranded cDNA, and this was followed by the ligation of barcoded TruSeq adapters. The products were purified and enriched via 10 cycles of PCR to create the final double-stranded cDNA library. To check library quality, we used the Bio-Rad QX200 Droplet Digital PCR EvaGreen Supermix System (Bio-Rad, USA) and a High Sensitivity DNA Analysis Kit (Agilent, USA). The prepared libraries were pooled for paired-end sequencing via Illumina HiSeq X Ten at Omics Drive Co. (International Plaza, Singapore) with 150 bp paired-ended sequence reads.

For RIP-seq, libraries were prepared with an Illumina TruSeq Stranded mRNA Sample Preparation Kit (Illumina, USA) according to the manufacturer’s protocol. Briefly, the purified RNA was fragmented and first-strand cDNA was synthesized using SuperScript III reverse transcriptase (Invitrogen, USA) with dNTP and random primers. Second-strand cDNA was generated using dUTP mix. The cDNA was modified and ligated to Accel-NGS 2S Indexed Adapters (Swift BioScience, USA) using an Accel-NGS 2S Plus DNA Library Kit (Swift BioScience, USA) according to the manufacturer’s instructions. 9N molecular identifiers (MIDs) on an i5 adapter were incorporated during ligation, allowing the identification of PCR duplicates. The final products were purified and enriched via 10 cycles of PCR using KAPA HiFi HotStart ReadyMix (Roche, Switzerland) and primer mix from a Swift Library kit to create the final double-stranded cDNA library. To check library quality, we used the Bio-Rad QX200 Droplet Digital PCR EvaGreen Supermix system (Bio-Rad, USA) and a High Sensitivity DNA Analysis Kit (Agilent, USA). The prepared libraries were pooled for paired-end sequencing via Illumina NovaSeq at Omics Drive Co. (International plaza, Singapore) with 150 bp paired-ended sequence reads.

### TRIBE analysis

Sequence reads were mapped to the *P. patens* genome V3.3 using the Bowtie2 and BLAT programs. Base positions with variants in each sample were analyzed using the PolymorphismCounter and NtFreqCounter functions in the RACKJ package (http://rackj.sourceforge.net/). We defined editing sites based on the following criteria: (i) the nucleotide at the site is covered by at least 20 reads; (ii) more than 80% of nucleotides at the site are adenosine in wild-type cells (thymine for reverse strand); (iii) more than 10% of nucleotides at the site have an adenosine-to-guanosine conversion (thymine-to-cytosine conversion for reverse strand) in *PTB2-HyperTRIBE*-overexpressing cells; (iv) the chi-squared *p* value on the site is calculated based on *G*-test log likelihood metric with a cutoff of *p* ≤ 1 × 10^−6^ for high-confidence editing sites (28). Editing sites detected in both replicates were further analyzed and annotated to different regions of genes based on the *P. patens* genome V3.3 using the MutationAnnotation function in the RACKJ package. The editing level is defined as the percentage of A-to-G conversion in *PTB2-HyperTRIBE*-overexpressing cells 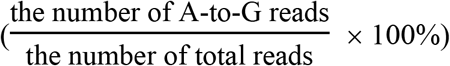.

### Validation of RBP targets in HyperTRIBE

DNase-treated total RNA was reverse transcribed into cDNA using Superscript III (Invitrogen, USA) according to the manufacturer’s protocol. PCR amplicons (∼250 bp regions upstream and downstream of editing sites) were cloned into *pBlueScript*. Approximately 27-30 clones were randomly selected and grown for plasmid extraction. The plasmids were subjected to Sanger sequencing, and the editing levels 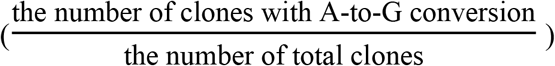 at each site within the PCR amplicon were calculated.

### Preparation of nuclear extract

For RNA immunoprecipitation, formaldehyde-fixed protonemata were ground into a fine powder in liquid nitrogen and resuspended in 30 ml extraction buffer 1 (400 mM sucrose, 10 mM Tris-HCl, pH 8.0, 10 mM MgCl_2_, 5 mM β-mercaptoethanol, 1 mM PMSF, 1× protease inhibitor cocktail, 50 U/ml SUPERaseIn RNase inhibitor) on ice for 5 min. The suspensions were filtered through six layers of Miracloth into a 50 ml Falcon tube. The filtrates were centrifuged at 3000 × *g* at 4°C for 5 min. The pellets (containing nuclei) were resuspended in 1 ml of extraction buffer 2 (250 mM sucrose, 10 mM Tris-HCl, pH 8.0, 10 mM MgCl_2_, 1% TX-100, 5 mM β-mercaptoethanol, 1 mM PMSF, 1× protease inhibitor cocktail, 50 U/ml SUPERaseIn RNase inhibitor), transferred to Eppendorf tubes, and centrifuged at 12,000 × *g* at 4°C for 5 min. The pellets were resuspended in 300 μl of extraction buffer 3 (1.7 M sucrose, 10 mM Tris-HCl, pH 8.0, 2 mM MgCl_2_, 0.15% TX-100, 5 mM β-mercaptoethanol, 1 mM PMSF, 1× protease inhibitor cocktail, 50 U/ml SUPERaseIn RNase inhibitor), layered on top of 600 μl extraction buffer 3 in Eppendorf tubes, and centrifuged at 16,000 × *g* at 4°C for 10 min. Finally, the pellets were resuspended in 300 μl nuclei lysis buffer (50 mM Tris-HCl, pH 8.0, 10 mM EDTA, pH8.0, 1% SDS, 1 mM PMSF, 1× protease inhibitor cocktail, 160 U/ml SUPERaseIn RNase inhibitor) and incubated on ice for 10 min for the RIP experiment.

### RNA immunoprecipitation

Protonemata (fresh weight ∼1.7 g) were fixed in 40 ml of 1% formaldehyde in TE buffer under a vacuum for 15 min. The process was stopped by adding 2 ml of 2.5 M glycine under a vacuum for 5 min. The protonemata were dried with tissue paper and frozen in liquid nitrogen. For each sample, 300 μl of nuclear extract was prepared and sonicated using a Bioruptor for 8 min at 4°C (power, high; 15 sec ON/15 sec OFF; 16 cycles). The extracts were centrifuged at 12,000 × *g* at 4°C for 1 min, and the supernatants were transferred to Eppendorf tubes. Total protein concentrations were measured using a Pierce BCA Protein Assay kit (Thermo Fisher Scientific, USA) to ensure that all samples had similar concentrations prior to immunoprecipitation. For each sample, 100 μl anti-cMyc (9E10)-conjugated magnetic beads were prepared by pre-blocking in 1 ml blocking buffer (16.7 mM Tris-HCl, pH 8.0, 1.2 mM EDTA, pH 8.0, 167 mM NaCl, 1.1% Triton X-100, 1 mM PMSF, 1× protease inhibitor cocktail, 160 U/ml SUPERaseIn RNase inhibitor, 1 mg/ml BSA, 0.5 mg/ml salmon sperm DNA) and washing once with 1 ml ChIP dilution buffer (16.7 mM Tris-HCl, pH 8.0, 1.2 mM EDTA, pH 8.0, 167 mM NaCl, 1.1% Triton X-100, 1 mM PMSF, 1× protease inhibitor cocktail, 160 U/ml SUPERaseIn RNase inhibitor). Each 1.2 mg extract sample was diluted 9-fold in ChIP dilution buffer, added to the anti-cMyc magnetic beads, and incubated on a rotator at 4°C overnight. After incubation, the beads were washed four times with 1 ml wash buffer (20 mM Tris-HCl, pH 8.0, 2 mM EDTA, pH 8.0, 300 mM NaCl, 1% Triton X-100, 0.1% SDS, 1 mM PMSF, 1× protease inhibitor cocktail, 40 U/ml SUPERaseIn RNase inhibitor) at 25°C for 5 min. A one-tenth volume aliquot of beads was prepared for immunoblotting to check for protein expression. The remaining nine-tenth volume of beads was used for RNA extraction. Briefly, the beads were treated with 1 mg/ml proteinase K at 50°C for 45 min and incubated at 95°C for 15 min for reverse crosslinking. The RNA was extracted using TRIzol reagent following the manufacturer’s protocol (Thermo Fisher Scientific, USA). To facilitate RNA precipitation, 2 μl of 5 mg/ml linear acrylamide and 35 μl of 3 M sodium acetate, pH 5.2, were added to the aqueous phase prior to precipitation in isopropanol. The RNA precipitants were treated with 100 μl of 0.04 U Turbo DNase I at 25°C for 30 min to remove DNA contamination and cleaned using an RNA Clean & Concentrator-5 kit following the manufacturer’s protocol (Zymo Research, USA). The RNA was eluted in 16 μl RNase-free water for subsequent experiments, such as RT-PCR and RIP-seq.

### RIP-seq analysis

Sequence reads were mapped to the *P. patens* genome V3.3 using the Bowtie2 and BLAT programs. The reads per kilobase per million mapped reads (RPKM) value for each gene was calculated using the RPKMComputer function in the RACKJ package and converted to transcripts per mission (TPM) values for subsequent statistical tests. To identify the enriched transcripts in the RIP-seq data, a Student’s *t*-test was performed by comparing the TPM values in three biological replicates between IP (*PTB2-5myc*) and mock (wild-type) samples. Transcripts that showed a cutoff *p* ≤ 0.01 and a 2-fold increase were defined as RIP enriched.

### Validation of RBP targets in RIP

RNA (3 μl) from the IP fraction in RIP was reverse transcribed into cDNA using Superscript III according to the manufacturer’s protocol (Invitrogen, USA). Fragments of ∼250 bp near the editing sites were PCR amplified and resolved by agarose gel electrophoresis.

### Data submission and accession numbers

RNA-seq and RIP-seq data have been submitted to the National Center for Biotechnology Information Sequence Read Archive database (http://www.ncbi.nlm.nih.gov/sra) with a BioProject accession number PRJNA735437. Gene information described in this article can be found in the Phytozome database of JGI (https://phytozome.jgi.doe.gov/pz/portal.html#!info?alias=Org_Ppatens) under the following gene locus numbers: *PpPTB2* (Pp3c16_18970).

## Acknowledgments

We thank Shu-Jen Chou and Ai-Ping Chen at the Genomic Technology Core Laboratory, Wen-Dar Lin at the Bioinformatics Core Laboratory of the Institute of Plant and Microbial Biology, Academia Sinica for technical assistance. We also thank Mitsuyasu Hasebe for sharing the pPGX vectors for β-Estradiol-Inducible system in *P. patens*.

## Author Contributions

Y.-L.C., H.-Y.H., and S.-L.T. designed the experiments; Y.-L.C. performed the experiments; H.-Y.H. and Y.-L.C. conducted RNA-seq data analysis; Y.-L.C. and S.-L.T. wrote the manuscript; S.-L.T. coordinated the project.

## Funding

This work was supported by a grant to S.-L.T. from Academia Sinica (Grant No. AS-108-TP-L01) and Ministry of Science and Technology (Grant No. MOST 109-2311-B-001 -029 -MY3).

## Figure legends

**Figure S1.**
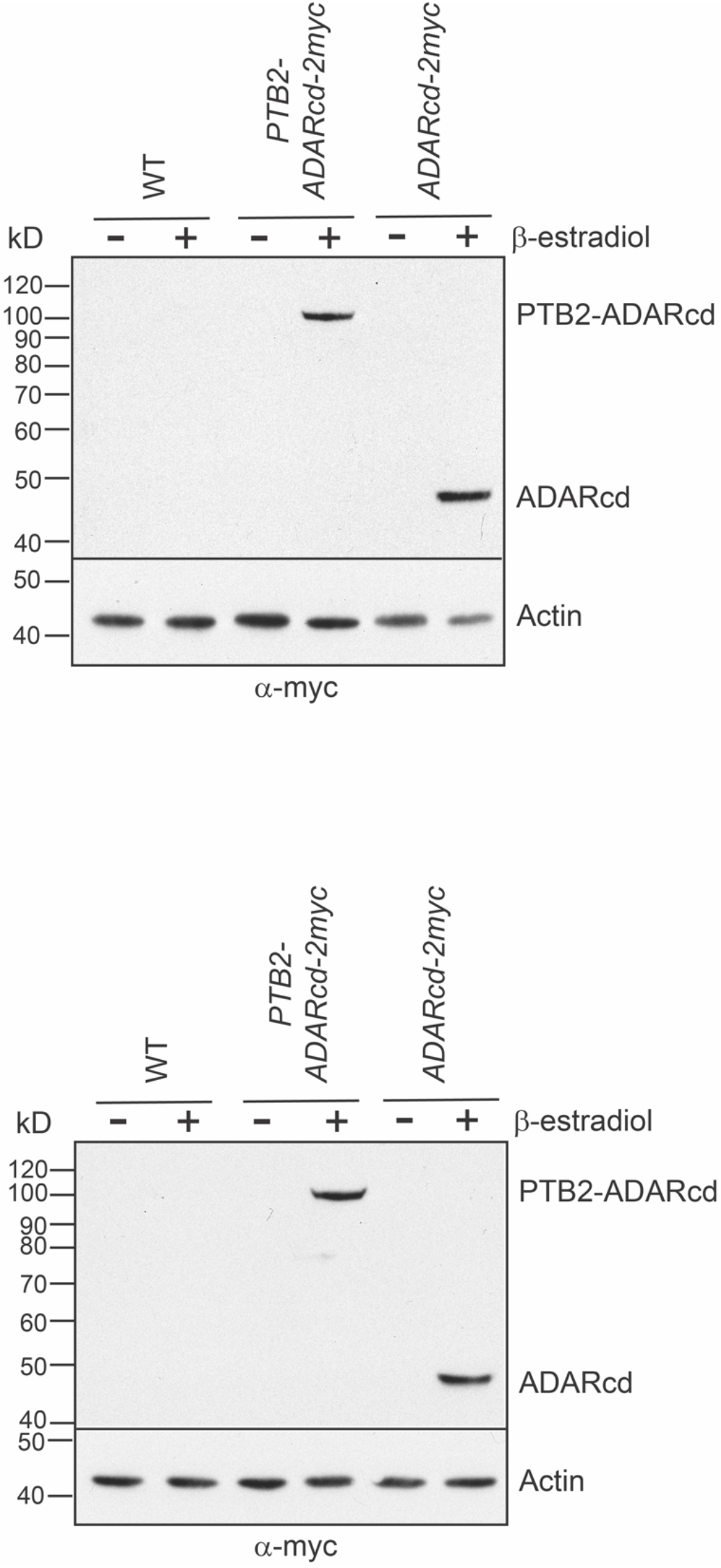
Immunoblotting of PTB2-HyperTRIBE with two biological replicates. Eight-day-old protonemata were treated with 10 μM β-estradiol for PTB2-HyperTRIBE induction. Total proteins were extracted from moss cells, resolved by SDS-PAGE, and transferred to a membrane for immunoblotting. Actin was used as the loading control.

**Figure S2.**
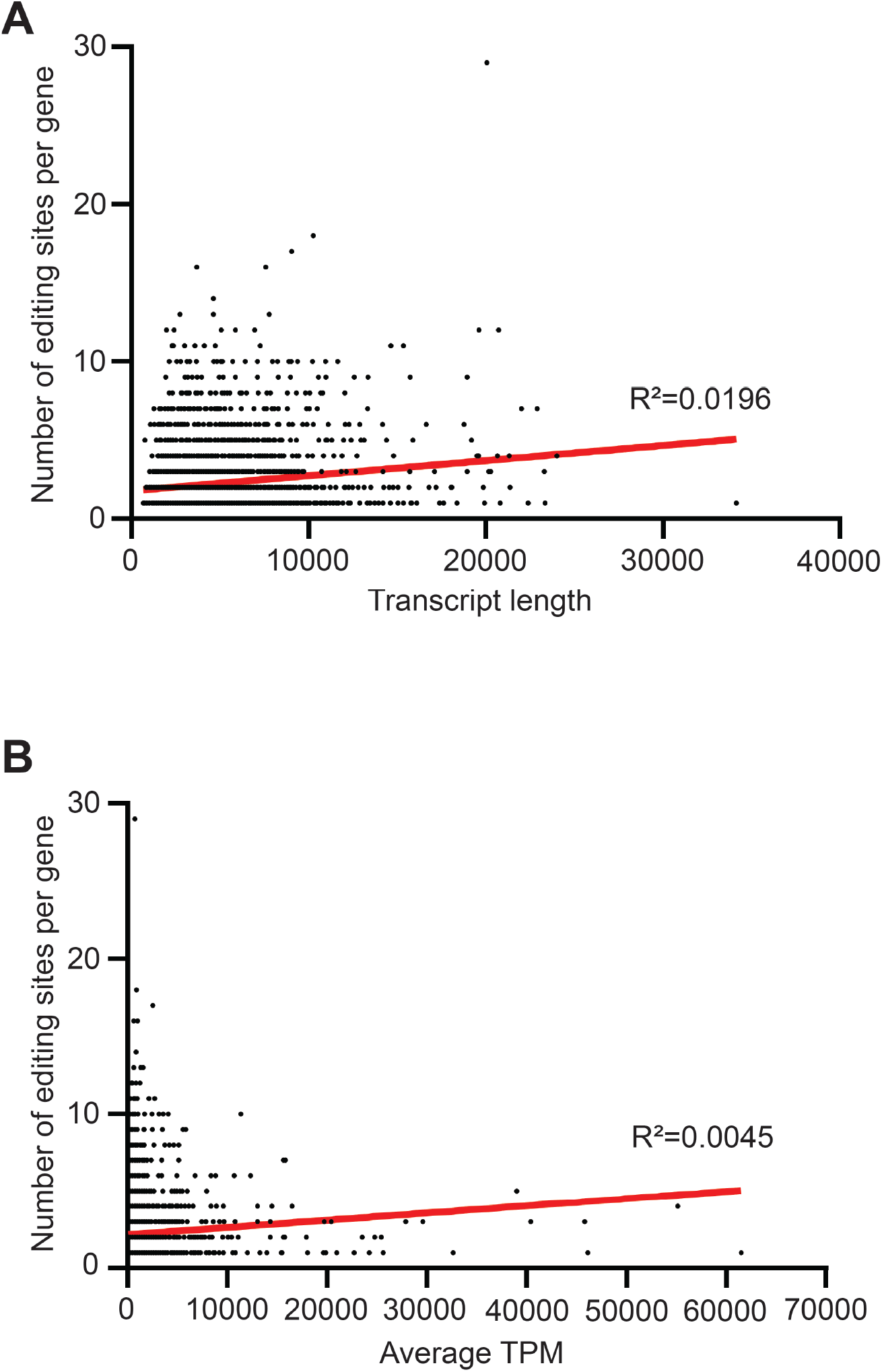
The number of editing sites does not increase proportionally in long or highly expressed transcripts. (A) Correlation between the number of editing sites and transcript length. (B) Correlation between the number of editing sites and transcript abundance.

## Notes

### Competing Interest Statement

The authors have declared no competing interest.

### Summary of Updates

figure arrangement and minor text updated

